# StructureDistiller: Structural relevance scoring increases resilience of contact maps to false positive predictions

**DOI:** 10.1101/697839

**Authors:** Sebastian Bittrich, Michael Schroeder, Dirk Labudde

## Abstract

Protein folding and structure prediction are two sides of the same coin. We propose contact maps and the related techniques of constraint-based structure reconstruction as unifying aspect of both processes. The presented Structural Relevance (SR) score quantifies the contribution of individual contacts and residues to structural integrity.

It is demonstrated that entries of a contact map are not equally relevant for structural integrity. Structure prediction methods should explicitly consider the most relevant contacts for optimal performance because they effectively double resilience toward false positively predicted contacts. Furthermore, knowledge of the most relevant contacts significantly increases reconstruction fidelity on sparse contact maps by 0.4 Å.

Protein folding is commonly characterized with spatial and temporal resolution: some residues are Early Folding while others are Highly Stable with respect to unfolding events. Using the proposed SR score, we demonstrate that folding initiation and structure stabilization are distinct processes.

---

Proteins are chains of amino acids which adopt complex, three-dimensional structures. This particular arrangement allows proteins to catalyze chemical reactions, transmit signals between cells, or recognize other molecules. The connection of protein sequence and structure is unclear and constitutes the protein folding problem. One promising technique to gain detailed insights into the process of protein folding (Figure 1) are pulse-labeling hydrogen-deuterium exchange (HDX) experiments [1, 2, 3]. In the process of protein folding, a denatured protein chain adopts a native, functional conformation. HDX allows to study the process with spatial and temporal resolution and folding events of particular residues can be related to particular time steps. Early Folding Residues (EFR, blue in Figure 1) initiate the formation of stable local structures starting from the denatured protein chain [1, 2, 4, 5]. In contrast, Highly Stable Residues (HSR, green in Figure 1) constitute regions in the native conformation [6] which are resilient to unfolding events (e.g. as natural phenomenon [7] or change in temperature or pH [8]). Both EFR and HSR are key aspects to understand the protein folding process [9, 3]; standardized data is provided by the Start2Fold database [10]. The defined-pathway model was proposed based on these observations. It considers protein folding to be a deterministic process where defined regions initiate the folding process and fragments assemble stepwise to form the native conformation [11, 2, 12] by establishing tertiary contacts [13, 14, 15, 2]. EFR constitute the folding nucleus and seem to determine the order in which certain sequence fragment fold. However, the relevance of EFR on the structural integrity of a protein structure is little explored. One reason is that it is currently not possible to assess the role of a contact or residue regarding the structural integrity of a protein; especially an *in silico* approach suitable for large-scale studies is needed to assess the relevance of EFR and HSR. Closely related to the protein folding problem are protein design and the prediction of structures from sequence [16].

**Figure 1:**
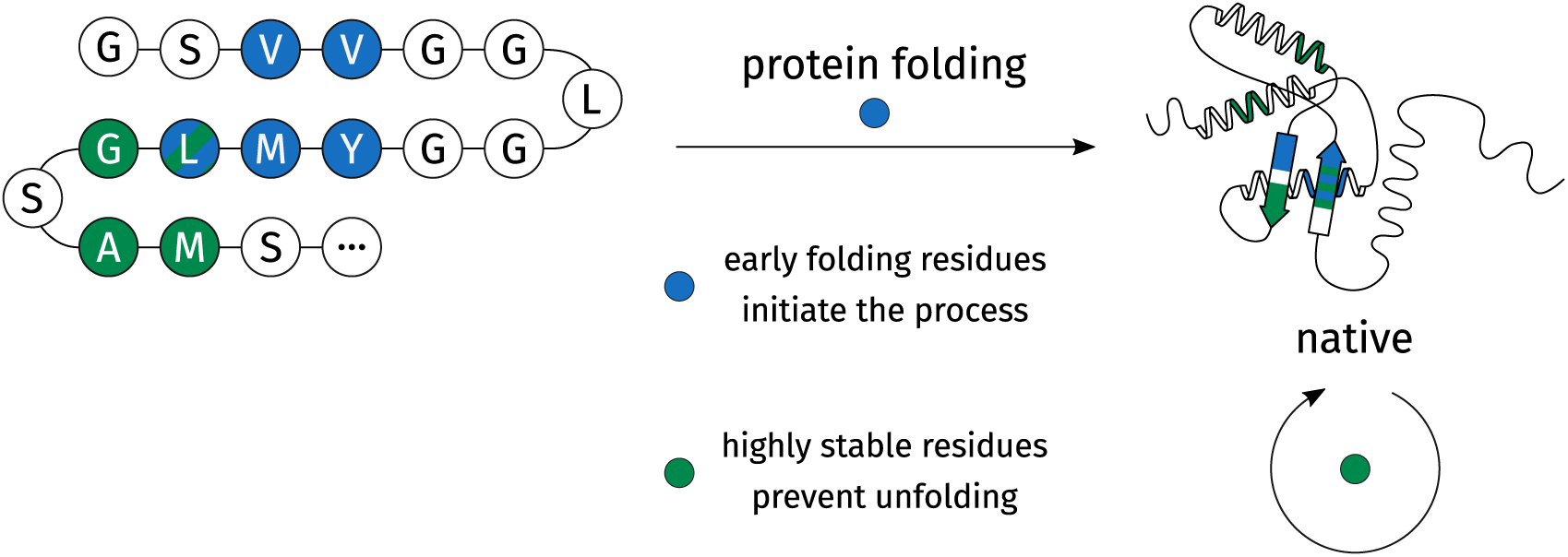
Studying protein folding by hydrogen-deuterium exchange. Most proteins adopt a native conformation autonomously in the process of protein folding [16, 17]. A small number of Early Folding Residues (EFR, depicted in blue) initiate the folding process as their surroundings change before that of other residues [3]. Analogously, folded proteins can be analyzed with respect to their stability. Highly Stable Residues (HSR, depicted in green) comprise regions which are particularly resilient to unfolding events [6].

Coevolution techniques [18, 19, 20, 21] propose an elegant approach to predict the structure of proteins from the abundance of sequences known today. For a given sequence, homologous sequences are retrieved and subsequently aligned via multiple sequence alignment (Figure 2a). Therein, some residues at defined sequence positions are conserved while others may change freely. A small number of residues are coupled to other positions: when one position changes the coupled position will change accordingly. This constraint implies the spatial proximity of both residues: even if they are separated at sequence level, they show a signal of coevolution because they are in contact at structure level (Figure 2b) [18]. The predicted contacts constitute a contact map (Figure 2c) which can be used as set of constraints for a subsequent structure reconstruction (Figure 2d). Conformations are sampled by a stochastic process in order to fulfill as many constraints as possible [22].

**Figure 2:**
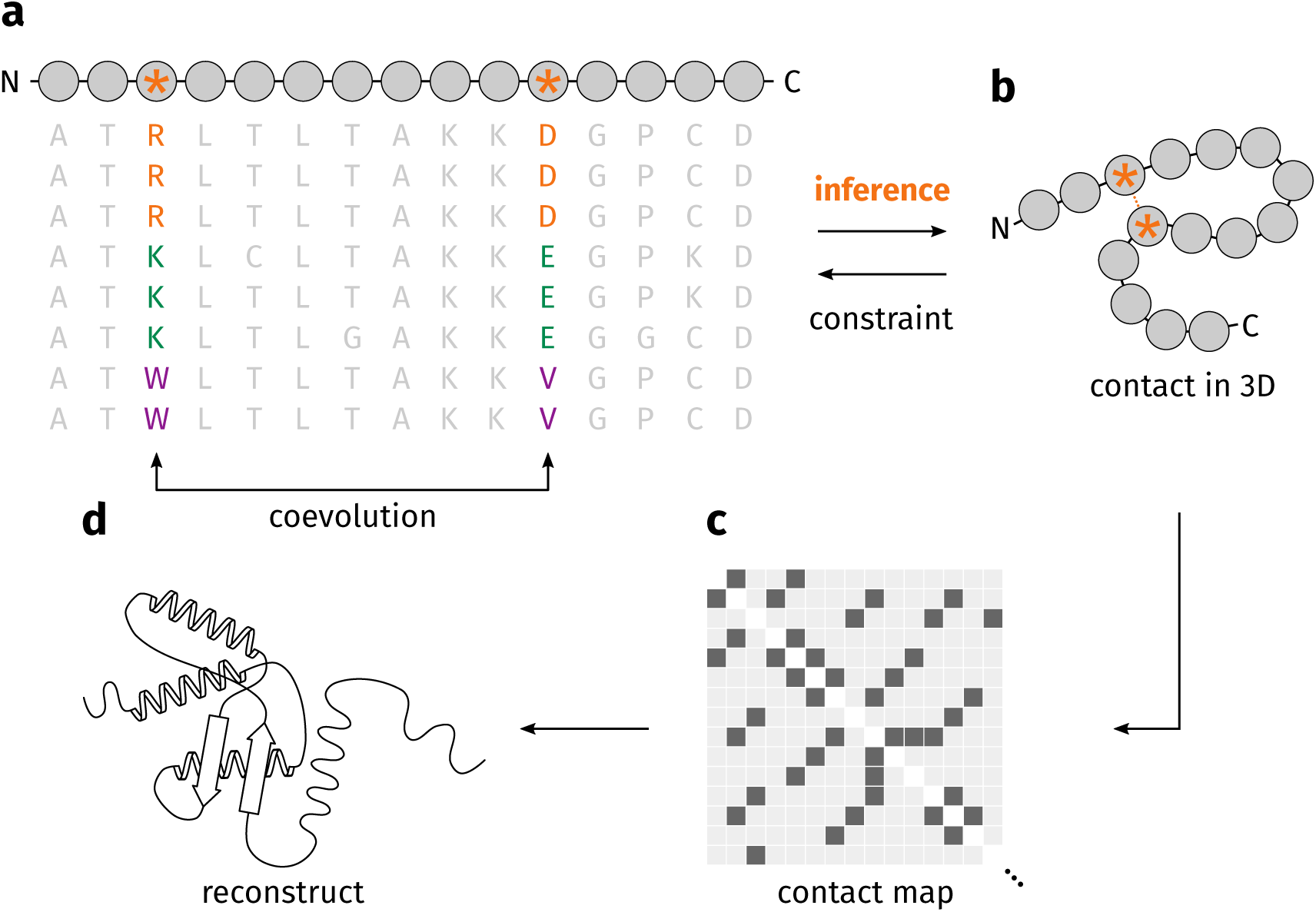
Protein structure prediction by coevolution techniques. (**a**) For a given sequence, homologous sequences can be used to create a multiple sequence alignment. Some positions coevolve (depicted by an orange asterisk): where for a change at one position a suitable change at the second position can be observed. (**b**) This connection at sequence level implies spatial proximity of both residues. (**c**) Coevolving residues can be represented by a contact map. (**d**) The predicted contacts are used as constraints of a subsequent structure reconstruction in order to find an optimal three-dimensional structure. Figure adapted from [18].

A contact map comprises the set of gathered constraints. Contact maps are matrices encompassing all pairs of sequence positions and usually contain a binary annotation whether two residues are in contact or not [23, 24]. They are used to design and train coevolution techniques and are also their output. Subsequently, these predicted contacts are used as constraints for reconstruction algorithms [22, 25, 26] in order to find conformations which fulfill the maximum number of constraints. Thus, coevolution techniques are capable of *ab initio* structure predictions which is not feasible by e.g. homology modeling approaches [27]. Predicted contacts used as constraints have also been demonstrated to speed-up molecular dynamics simulations by allowing for faster convergence [28].

The success of coevolution techniques continues to revolutionize structural biology [29, 19] and spawned a comprehensive ecosystem of related methods revolving around contact maps. The recent iteration of the CASP experiment emphasizes the coming of age of contact prediction and the improvement of *ab initio* protein folding protocols [30, 31, 32]. Dedicated methods for the visualization and interpretation of contact maps were created [33, 34, 35]. Quality assessment of the predicted contacts becomes increasingly important as well. False positive predictions (i.e. contacts not observed in the native structure of a protein) are common. They have detrimental effects on the usefulness of contact maps [23, 24]. Peculiarly, such false positive predictions are difficult to spot [18] and in reconstruction they impair the feasibility of all other contacts [36]. Thus, dedicated methods were designed to validate contact maps [35, 37]. Other studies [24] tried to elucidate the optimal contact definition by assessing its influence on reconstruction performance. Commonly, contacts stabilizing secondary structure elements (i.e. residues separated by less than six positions on sequence level) are ignored in the context of contact maps [38]. The range of the remaining contacts are considered short (sequence separation of 6–11), medium (12–23), or long (>23) [39].

Contact maps do not only contain the information needed for protein structure prediction, but they also are potential tools to describe the fundamentals of protein folding. In 2007, Chen et al. [40] pioneered the search for the most relevant contacts of a contact map and wanted to determine the minimal set of contacts which captures the fold of a protein. Therefore, they represented proteins by contact maps and selected random subsets with varying coverage. These subsets were then used as constraints in a structure reconstruction algorithm, the result was aligned to the native structure, and its fidelity was assessed by the root-mean square deviation (RMSD). As the number of constraints increased (i.e. more contacts of the native contact map are considered), the RMSD decreased because the reconstructs resembled the native structure increasingly well. A reconstruction is considered successful when the RMSD to the native structure is below a certain threshold and likely to resemble the correct fold [23, 41, 40]: in our study, we consider 4.0 Å as threshold. Good reconstructions have been shown to depend on a delicate balance of sequentially neighbored and sequentially separated contacts [40]. Sathyapriya et al. [41] extended the study of Chen et al. and coined the term *structural essence* for the minimal set of fold defining contacts. They demonstrated that 8% of all contacts allow for the reconstruction of the correct fold of a protein because most information in a contact map is redundant. Furthermore, a rational selection of contacts can outperform a random selection of equally many contacts with respect to reconstruction quality. However, such a configuration is difficult to compose [41]. Duarte et al. showed that consideration of all contacts leads to reconstruction qualities around 2 Å [24].

The annotation of EFR and HSR provided by the Start2Fold database [10] is valuable information to understand the protein folding problem and has also implications for the prediction of protein structures. Contact maps are the cornerstone of contemporary structure prediction methods. The surrounding ecosystem of reconstruction algorithms may elucidate the protein folding process by pinpointing the most important contacts for structural integrity. Additionally, the relevance of EFR and HSR in the context of protein structure prediction provides qualitative insights. Several studies identified a small number of key residues for the *in vitro* folding process. Is the same true for *in silico* folding: are some constraints more important than others? Are positions featuring evolutionary couplings crucial for reconstructions? For a long time, *in silico* folding simulations improved the understanding of the protein folding process [42, 43], potentially contact maps provide an even more tangible connection of both aspects. To address these questions, we propose the Structural Relevance (SR) score which captures the performance increase that considering an individual contact or residue for a *in silico* reconstruction process provides.

## Results

A subset of proteins from the Start2Fold database [10] were analyzed. The folding and stability characteristics of the corresponding proteins have been determined by HDX experiments [8, 10, 44] and these properties may relate to the most relevant contacts of a contact map and constitute a direct connection of protein folding *in vivo* and structure prediction *in silico*. We only considered entries for which both EFR and HSR were annotated, totalling 30 proteins. Individual contacts cannot be directly assessed regarding their SR score (i.e. how much does knowledge of this contact improve reconstructions) because a single contact will never yield a meaningful reconstruct, instead they depend on a set of other contacts [40, 41]. In order to quantify the structural relevance of individual contacts, they have to be disentangled from all other contacts mandatory for a meaningful reconstruction in the first place (see method section). The reconstruction error describes the dissimilarity of each reconstruct with respect to the native structure [40, 41].

For a basic assessment of reconstruction quality, all proteins in the dataset were reduced to a contact map representation and random subsets with varying coverage were used to reconstruct all proteins (Figure 3). With increasing number of considered contacts, the reconstruction error decreases. The reconstruction process using more contacts becomes more robust as the distributions decrease in variance. See detailed results and the list of Start2Fold entry identifiers in Supplementary Figure 1. At 30% coverage of all native contacts the yielded reconstructs resemble the fold of the native structure and are also sensitive to the removal or addition of individual contacts. The reconstruction error approaches 2 Å when all contacts are used as described in literature [41].

**Figure 3:**
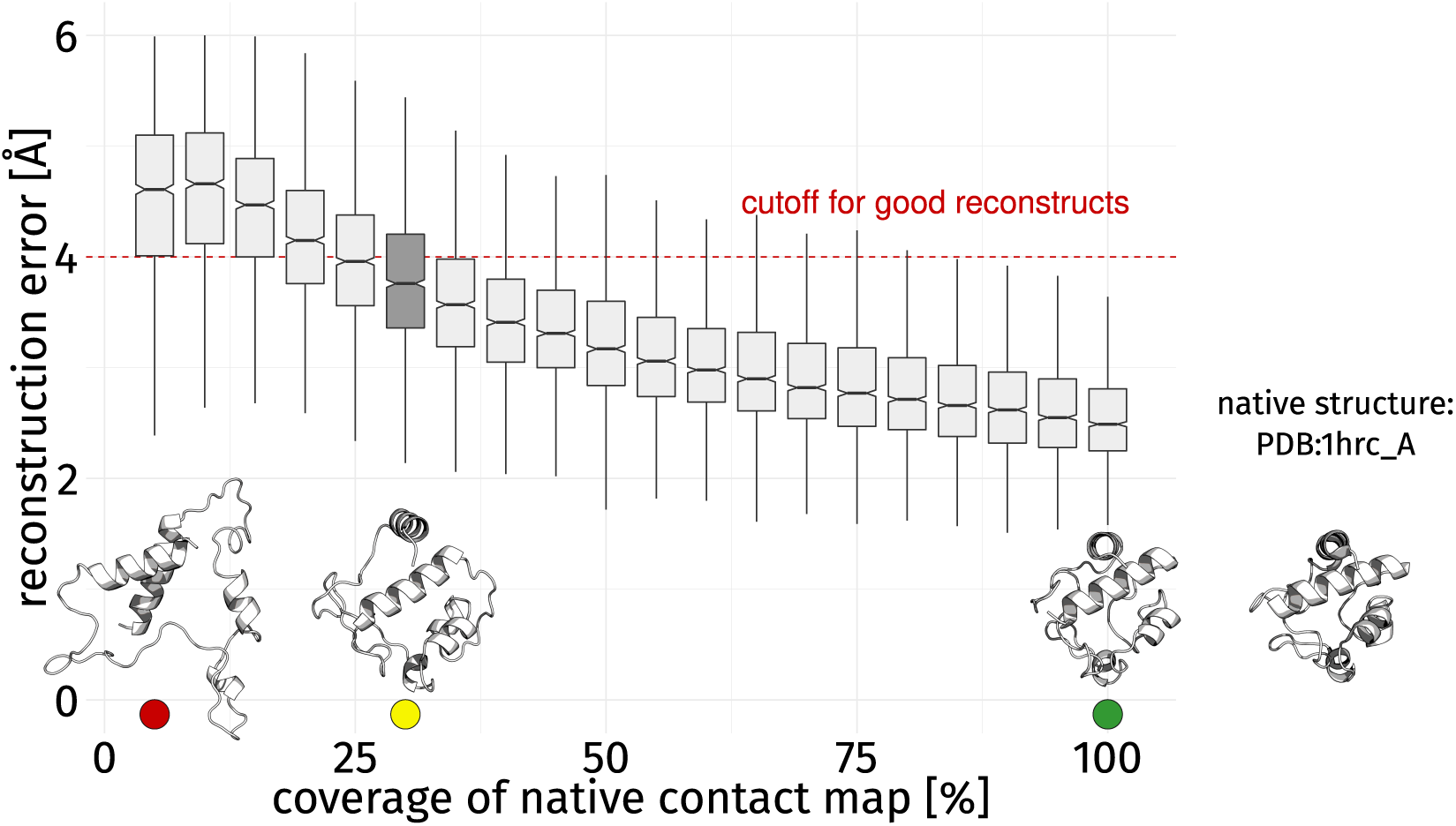
Reconstruction error by percentage of contacts. When more contacts are considered, the average reconstruction error decreases [40] and the same is true for the variance of each bin. For the assessment of the SR of contacts, 30% of all native contacts (box plot filled dark gray) were chosen as compromise because it ensures reconstructs of average quality while the corresponding contact maps are still sensitive to the removal or addition of individual contacts (as indicated by a big shift in reconstruction error with respect to the neighboring bins). Renderings of four structures are provided to make the influence of the coverage of the native contact map more tangible. They resemble knowledge of 5%, 30%, and 100% of all native contacts as well as the native structure (PDB:1hrc A, isolated on the right).

### The Structural Relevance of individual contacts and residues

The SR score of 5,173 contacts was computed by our StructureDistiller algorithm (see method section). The outputted score captures the average performance increase in Å, when a particular contact is considered for the reconstruction process compared to a reconstruction without knowledge of this contact (ΔRMSD). Positive SR scores indicate contacts which favorably contribute to reconstruction fidelity, whereas negative scores indicate native contacts which hinder or at least not substantially improve the process. The removal of an individual contact results in (negative) change in SR by 0.012 ± 0.253 Å (throughout the manuscript the standard error is given). In contrast, the addition of a contact leads to an increase by 0.022 ± 0.253 Å. Most contacts contribute positively to reconstruction performance. Only a small number of contacts is of high SR with similar tendencies shown by studies on contact maps [40, 41] as well as protein folding in general [45], where good reconstructions as well as correctly folded protein structures depend on a small number of key contacts. The high variance of the SR scores is the result of both the contact map sampling as well as the reconstruction routine [22] being stochastic processes. Both operations are performed with ten-fold redundancy to limit this issue. The presented SR scores are the average values over all redundant runs.

We used several features (Table 1) to describe contacts in the dataset in more detail and assess their relation to the SR score. Therefore, residue contacts are distinguished according to their sequence separation [39]. Short contacts (6–11) exhibit a significant decrease in the SR score. In contrast, long contacts (>23) of sequentially highly separated residues are more common and feature increased SR scores. The change is insignificant for contacts of medium range. Non-covalent interactions such as hydrogen bonds and hydrophobic interactions are annotated between residues [46] and contacts which are backed by either contact type are considered. Both are associated with a significant change whereby the presence of hydrogen bonds decreases the SR and hydrophobic interactions increase the SR of a contact.

**Table 1:** Contact-level features influencing the SR (ΔRMSD) score.

| Table 1: Contact-level features influencing the SR (ΔRMSD) score. | | | | | | |
| --- | --- | --- | --- | --- | --- | --- |
| feature | present | $n$ | $\mu_{\text{SR}}$ [pm] | $\sigma_{\text{SR}}$ [pm] | trend | $p$-value |
| short contact (6–11) | yes | 1,120 | 1.4 | 8.1 | ↓ | 0.025 |
|  | no | 4,053 | 2.2 | 8.6 |  |  |
| medium contact (12–23) | yes | 1,271 | 1.8 | 8.6 | – | 0.161 |
|  | no | 3,902 | 2.1 | 8.5 |  |  |
| long contact (> 23) | yes | 2,782 | 2.4 | 8.6 | ↑ | 0.002 |
|  | no | 2,391 | 1.6 | 8.4 |  |  |
| hydrogen bond | yes | 563 | 1.1 | 8.3 | ↓ | 0.018 |
|  | no | 4,610 | 2.1 | 8.5 |  |  |
| hydrophobic interaction | yes | 541 | 2.9 | 9.5 | ↑ | <0.001 |
|  | no | 4,632 | 1.9 | 8.4 |  |  |
| evolutionarily coupling | yes | 1,461 | 2.2 | 8.4 | – | 0.203 |
|  | no | 3,246 | 1.8 | 8.7 |  |  |
| top-scoring coupling | no | 3,687 | 1.8 | 8.7 | – | 0.059 |
|  | yes | 1,020 | 2.4 | 8.3 |  |  |

Contact length refers to the sequence separation of the contact [39]. Hydrogen bond and hydrophobic interaction refers to contacts for which the respective interaction type was observed [46]. Evolutionary couplings by direct coupling analysis [18, 49], for some proteins no data could be computed. Top-scoring couplings are the first $0.4L$ contacts sorted by their coupling rank. $n$ describes the number of observations, $\mu$ the corresponding average, and $\sigma$ the respective standard deviation. The trend is given, i.e. does presence of this feature decrease (↓) or increase (↑) the SR scores. Insignificant change is represented by a dash (–).

Previously it has been shown that contacts within as well as between secondary structure elements are required for optimal reconstruction performance [40, 41]. Commonly, reconstructions only consider residue pairs at least six positions apart on sequence level [39], though there are cases where the usually ignored contacts may contribute valuable information on the structure of loops [47]. Non-covalent interactions have a significant effect on the SR score of a contact. Hydrogen bonds occur between backbone atoms of amino acids where they define and stabilize secondary structure elements. Some amino acids such as serine or threonine feature polar side chains which allow them to engage more flexibly in this type of non-covalent interaction. The importance of hydrogen bonds furnished by side chains for protein folding and stability has been shown [48, 16]. Hydrogen bonds may feature lower SR scores because of their propensity to occur between polar amino acids at positions exposed to the solvent. In contrast, hydrophobic interactions primarily occur in the buried hydrophobic core of a protein where they are surrounded by many other residues which reduces the degree of freedom. Especially, the importance of tertiary contacts furnished by hydrophobic interactions has been shown [14, 5]. Such interactions provide information on the correct assembly of distant parts of the protein and, thus, are relevant for structural integrity both during protein folding and in structure prediction.

Regarding contact prediction methods, evolutionary couplings [18, 49] show no significant association to the SR score. However, a slight increase in SR can be observed, when two positions are evolutionarily coupled. A selection of the 0.4*L* top-scoring contacts (*L* refers to the sequence length) results in a more substantial, though still insignificant, change in SR. Many predicted couplings are not actually present in the native contact map due to the strict distance cutoff. Also, potential false positive predictions by the direct coupling analysis are not evaluated, which can be expected to have a negative effect on reconstruction quality [23].

At residue level, a set of features was evaluated with the same reasoning (Table 2). Residues in loop regions have significantly lower SR than those in *α*-helices and *β*-strands. For secondary structure elements, backbone angles and hydrogen bonding patterns are used as additional constraints during reconstruction [22] which may explain an overall performance increase. The previous association of hydrophobic interactions and SR score may be explained by a bias for buried residues; however, no significant association is observed at residue level. The annotation of EFR does not influence SR scores significantly, while the opposite is true for HSR (see below). Functional residues may not be of high SR, because binding sites tend to be exposed to the solvent and commonly have unfavorable conformations [50]. Again evolutionary couplings [18, 49] do not lead to increased SR, probably because most residues feature at least one predicted coupling. Filtering for the 0.4*L* top-scoring positions (i.e. regarding their cumulative coupling strength) does not lead to a significant change either.

**Table 2:** Residue-level features influencing the average SR (ΔRMSD) score.

| Table 2: **Residue-level features influencing the average SR (ΔRMSD) score.** |  |  |  |  |  |  |
| --- | --- | --- | --- | --- | --- | --- |
| **feature** | **present** | **$n$** | **$\mu_{\text{SR}}$ [pm]** | **$\sigma_{\text{SR}}$ [pm]** | **trend** | **$p$-value** |
| early folding | yes | 414 | 2.6 | 6.1 | — | 0.543 |
|  | no | 2,115 | 2.5 | 6.2 |  |  |
| highly stable | yes | 688 | 3.0 | 6.2 | ↑ | < 0.001 |
|  | no | 1,731 | 2.1 | 6.1 |  |  |
| functional | yes | 119 | 2.8 | 6.2 | — | 0.919 |
|  | no | 2,078 | 2.6 | 5.9 |  |  |
| coil | yes | 996 | 1.9 | 6.4 | ↓ | < 0.001 |
|  | no | 1,533 | 2.9 | 6.0 |  |  |
| buried | yes | 1,105 | 2.7 | 5.5 | — | 0.075 |
|  | no | 1,424 | 2.4 | 6.7 |  |  |
| evolutionarily coupled | yes | 1,975 | 2.6 | 6.2 | — | 0.754 |
|  | no | 503 | 2.6 | 6.4 |  |  |
| top-scoring coupled | yes | 1,117 | 2.6 | 5.7 | — | 0.492 |
|  | no | 1,361 | 2.5 | 6.6 |  |  |

Residues in coil regions and residues buried according to their relative accessible surface area were evaluated. Residues were assessed regarding their early folding and highly stable characteristics [10]. Annotation of functional residues from UniProt [67]. Considers evolutionary couplings and the $0.4L$ top-scoring positions according to the cumulative coupling strength [18, 49]. $n$ describes the number of observations, $\mu$ the corresponding average SR, and $\sigma$ the respective standard deviation. The trend is given, i.e. does presence of this feature decrease (↓) or increase (↑) the SR scores. Insignificant change is represented by a dash (—).

### Most relevant contacts increase reconstruction performance

The subset of contacts with high SR scores should lead to good reconstructs when combined. To test this hypothesis, proteins were reconstructed using various subset selection strategies equal to 30% of all native contacts (Figure 4). A baseline is obtained by selecting 30% of the contacts randomly (gray). Rational selections are based on sorting all contacts in a protein by its SR scores. The 30% top-scoring contacts represent the most relevant contacts (green). The bottom 30% represent the least relevant contacts (red). Other interesting aspects are contact distance and type: therefore short (6–11), long (>23) contacts, hydrogen bonds, and hydrophobic interactions were assessed (Supplementary Figure 2).

**Figure 4:**
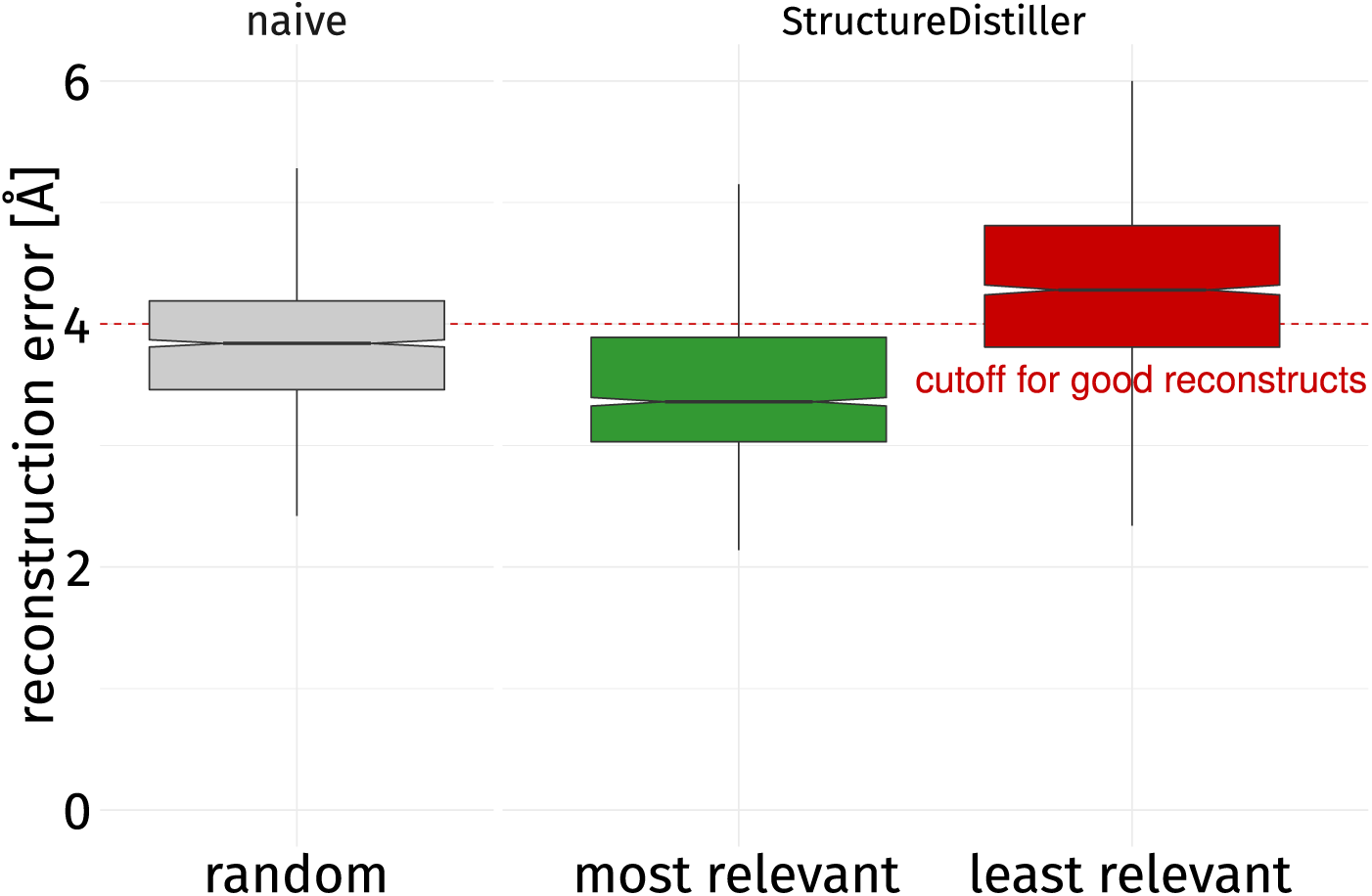
Impact on reconstruction performance by strategy. Three strategies were used to reconstruct structures of the dataset using a number of constraints equal to 30% of contacts in the native map. A random selection of contacts (gray), the most relevant ones by SR score (green), and the least relevant ones (red). The most relevant contacts yield the lowest reconstruction error when combined. This configuration outperforms a random selection of contacts significantly (*p*-value: <0.001). Previous studies [40, 41] have shown the difficulties in finding combinations of contacts yielding better reconstructs than a random selection. Using the least relevant contacts results in an increased error compared to the random selection (*p*-value: <0.001). When only a subset of all entries of a contact map can be considered (as it is commonly the case [35] and reasonable for efficiency [41]), the subset of contacts chosen is crucial for reconstruction performance.

The RMSD is used to quantify the fidelity of a reconstruct by aligning it to the native structure – high reconstruction errors occur for bad reconstructs. A random selection of 30% of contacts achieves 3.839 ± 0.599 Å. A combination of contacts by the most relevant strategy significantly outperforms the random strategy with an average reconstruction error of 3.479 ± 0.625 Å. Consideration of the least relevant contacts results in an increase in reconstruction error to 4.311 ± 0.687 Å.

Chen et al. assumed that no rational selection of contacts can surpass a random selection in terms of reconstruction fidelity [40]. Later, Sathyapriya and coworkers [41] provided an algorithm capable of doing just that. It is especially remarkable that their approach merely evaluates which neighborhood is shared by a pair of residues. The main aspect of their algorithm is the selection of non-redundant contacts which can provide the maximum amount of information for a reconstruction when combined. The selection of the most relevant contacts as determined by StructureDistiller constitutes a different approach to compose a set of contacts which allow for better reconstructs than a random selection. Of all native contacts two selections can be readily made. One is significantly better suited for reconstruction purposes than a random selection and whereas the other one performs significantly worse. It is also remarkable that a combination of long contacts performs significantly worse than the negated selection (Supplementary Figure 2), despite individual long contacts exhibiting high SR scores (Table 1). This emphasizes the context-specificity of individual contacts and substantiates the previous findings [40], wherein both short and long contacts are needed for good reconstructions.

### Increased resilience to false positive predictions

The sensitivity of a contact map to false positive contacts has been discussed before – even a small number of such contacts not present in the native structure is detrimental to reconstruction performance [23]. As shown in the previous section, contacts with high SR allow for better reconstructs. Interestingly, the selection of the most relevant contacts also can compensate the moderate introduction of false positive contacts (Figure 5). The selection of the most relevant contacts performs significantly better than a random selection in all considered cases. The introduction of false positive contacts quickly leads to reconstructions with errors above 4 Å as larger fractions of false positive contacts dilute the correct information captured by native contacts. When more than 7% false positive contacts are introduced to the most relevant selection, the majority of reconstructions is of bad quality. When 30% of all contacts are selected randomly, only 3% false positive contacts can be introduced before the error exceeds the threshold of 4 Å. The consideration of the most relevant contacts buffers the negative influence of false positive predictions (Table 3): median performance is comparable between reconstructions based on a random selection without false positive contacts and the selection of the best contacts diluted by 6% of false positive contacts.

**Figure 5:**
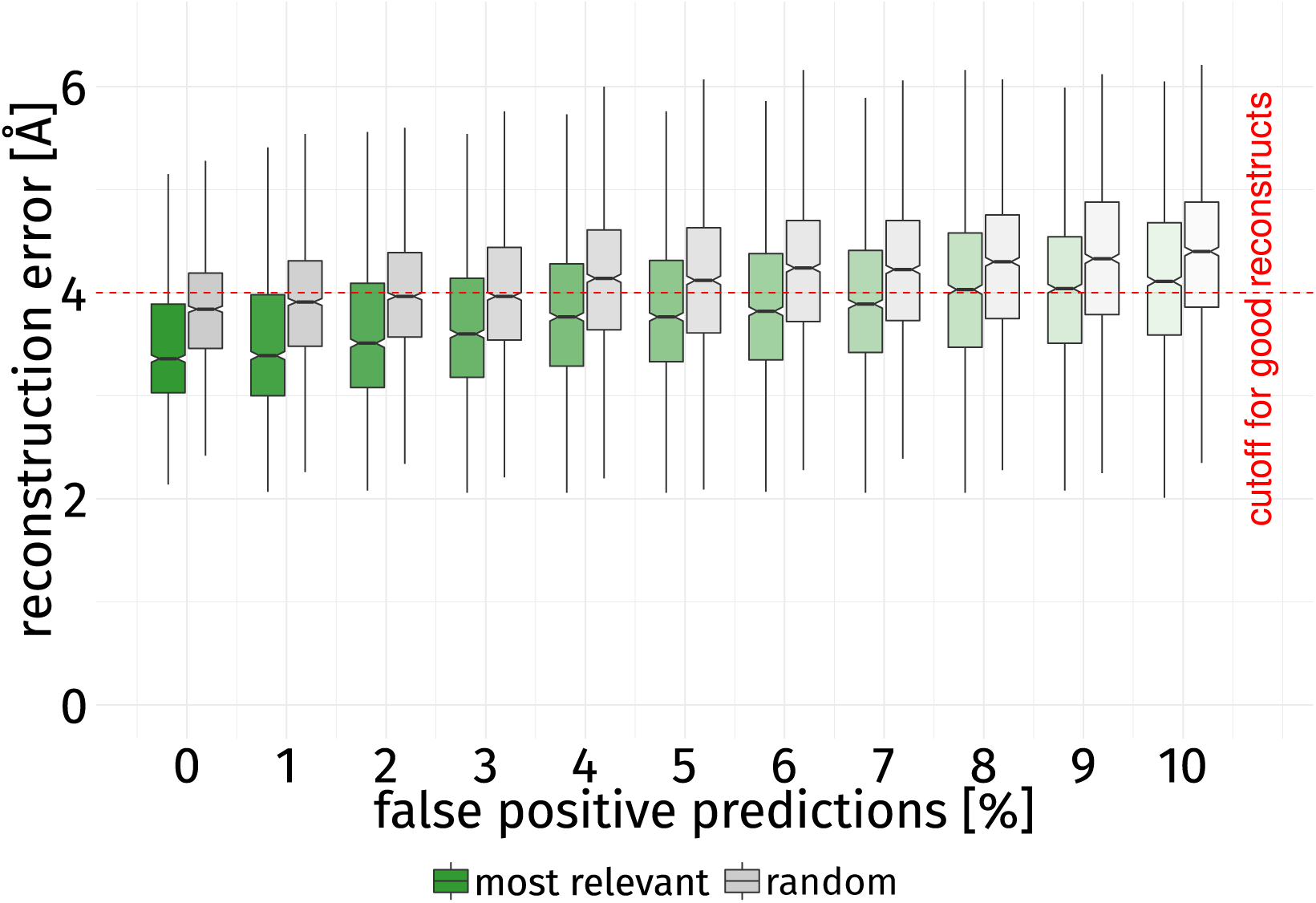
Influence of false positive contacts. The reconstruction error is given of for 30% of all contacts in the most relevant (green) and random (gray) bins with an increasing fraction of false positive contacts. In all cases, the most relevant contacts perform significantly better than a random selection when it comes to compensating false positive contacts (*p*-value <0.001). E.g., the median performance of a random selection without false positive contacts is comparable to that of the best selection with 6% false positive contacts. When more than 3% false positive contacts are introduced into the random selection, the error of the majority of reconstructions lies above 4 Å, whereas the best selection can compensate more than double the number of false positive contacts before surpassing this threshold. Knowledge of the most relevant contacts as quantified by the StructureDistiller algorithm thus increases the resilience to false positive contacts as well as the overall reconstruction performance.

**Table 3:** Reconstruction error introduced by false positive contacts.

| **false positive contacts [%]** | **$\mu_{\text{best}}$** | **$\tilde{x}_{\text{best}}$** | **$\mu_{\text{random}}$** | **$\tilde{x}_{\text{random}}$** |
| --- | --- | --- | --- | --- |
| 0 | 3.479 | 3.360 | 3.839 | 3.840 |
| 1 | 3.498 | 3.390 | 3.903 | 3.910 |
| 2 | 3.598 | 3.510 | 3.971 | 3.965 |
| 3 | 3.665 | 3.600 | 3.996 | 3.965 |
| 4 | 3.808 | 3.765 | 4.135 | 4.140 |
| 5 | 3.840 | 3.765 | 4.117 | 4.120 |
| 6 | 3.882 | 3.820 | 4.211 | 4.240 |
| 7 | 3.931 | 3.890 | 4.229 | 4.225 |
| 8 | 4.028 | 4.030 | 4.256 | 4.300 |
| 9 | 4.038 | 4.040 | 4.292 | 4.330 |
| 10 | 4.140 | 4.110 | 4.354 | 4.400 |

For increasing rates of false positive contacts the reconstruction performance using 30% of the native contacts are given. $\mu_{\text{best}}$ refers to the average performance using the most relevant contacts, $\mu_{\text{random}}$ to that using a random selection of contacts. $\tilde{x}$ describes the median of the corresponding population. In all cases, the performance of the best bin is significantly better than that of a random selection.

Since even those native contacts can hinder reconstruction (as indicated by negative SR scores), it becomes evident that the correct ranking of contacts [22, 37] has a serious influence on reconstruction quality and should be considered for the design and training of contact prediction techniques. The insignificant association of evolutionary couplings and SR scores suggests that the most relevant contacts may not be easy to predict but can contribute significantly more information needed for the successful reconstruction of a protein.

### Analysis of Early Folding and Highly Stable Residues

A direct connection to particular folding and stability characteristics is provided by the annotation of EFR which initiate and guide the folding process. However, according to the SR score we observe no change for EFR (Table 2). Contacts of HSR exhibit a significant increase in SR compared to unstable contacts. It is remarkable that contacts of EFR show no increase in SR despite their presumed role for the protein folding process [44, 4]. A possible interpretation is that EFR primarily define stable, local structures [44, 4] due to their occurrence in sequence regions associated to high backbone rigidity. They form defined sequence regions with fewer possible backbone conformations and produce pivotal secondary structure elements. Therefore, EFR define the folding nucleus of a protein and sequentially encode the ordered secondary structure elements formed first. However the obtained SR scores suggest that crucial contacts between these secondary structure elements may be mediated by other residues which are not necessarily EFR themselves, but may occur in secondary structure elements containing EFR [1].

Another aspect of the experimental data by Pancsa et al. is the annotation of residues which are strongly protected in stability measurements [10]. Such residues occur in ordered secondary structure elements and their contacts are beneficial to reconstruction performance. Rather than initiation the formation of the native structure (like EFR), HSR seem to manifest the native conformation. The differences in SR scores between EFR and HSR imply that two distinct process are realized by these two distinct sets of residues.

The defined-pathway model [11, 2, 12] describes protein folding as a deterministic, hierarchic process. EFR occur in regions which autonomously fold first relative to the rest of a protein. Furthermore, this tendency does not depend on tertiary contacts in a protein structure, but is rather the direct consequence of the local sequence composition [1, 10, 44]. These stable, local structures may be secondary structure elements [3] or larger autonomously folding units also referred to as foldons [2]. In a stepwise process, such local structures will subsequently establish tertiary contacts and assemble the native conformation of a protein [15, 51, 2]. The employed reconstruction method directly considers secondary structure elements, which are used to derive additional constraints. Therefore, most secondary structure elements should be represented successfully which may explain why we observe long contacts to be particularly important for structural integrity. It is also reasonable that the SR score of a contact increases with the distance at the sequence level: potentially, such constraints do not only enforce the correct placement of both residues but also have an indirect positive impact on the correct conformation of all residues in between.

### Disruption to cytochrome c induces molten globule state

Ground truth on the structural importance of individual contacts is difficult to find – as a case study cytochrome c is used, which is a member of the dataset. Cytochrome c (Figure 6) contains two Ω-loops which are stabilized by a hydrogen bond between HIS-26 and PRO-The importance of this contact has been shown as disruptions induce a molten globule state [52, 53]. Particularized folding studies [8] have also identified the N- and C-terminal helices as foldons, i.e. autonomously folding units which initiate and guide the folding process. Besides that, wide parts of the structure are constituted of coil regions and fixate a heme ligand, thus potentially exhibiting increased structural flexibility.

**Figure 6:**
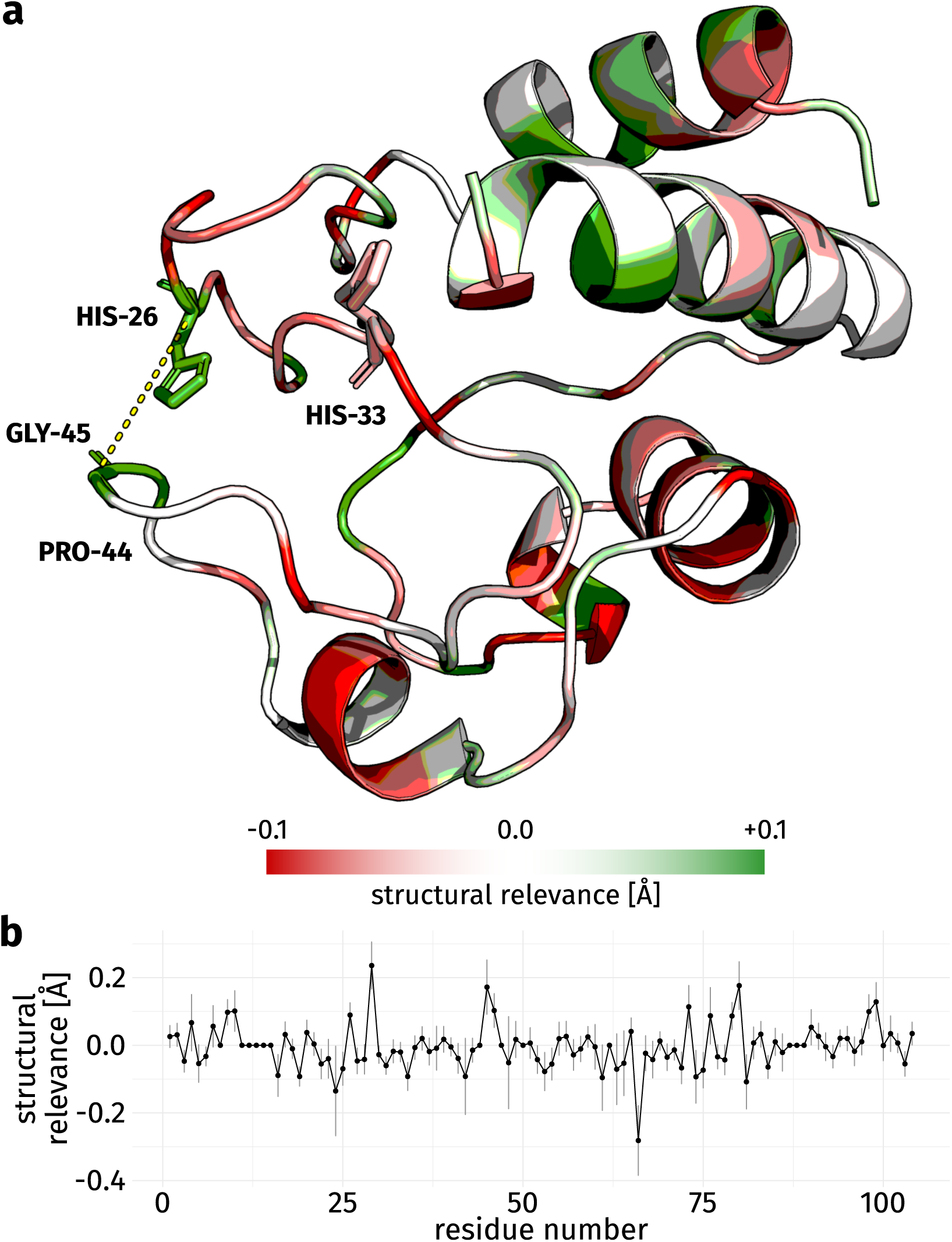
Cytochrome c (PDB:1hrc A) colored by Structural Relevance (ΔRMSD). (**a**) Residues with high SR scores are depicted in green, those with negative SR are rendered in red. For gray residues no contacts were observed and no SR scores are reported. It has been shown in experiment that disruptions to the hydrogen bond between HIS-26 and PRO-44 will induce a molten globule state when the association between both Ω-loops is lost [52, 53]. StructureDistiller reports high SR for HIS-26, GLY-45, and the contact both share (yellow dashed line), though no direct contact is detected between HIS-26 and PRO-44 due to strict distance threshold of the employed contact definition. HIS-33 has been described as variable position lacking any structurally relevant contacts [52] and this observation is manifested in the low SR score of this residue. The N- and C-terminal helices exhibit high SR, especially for residues which constitute their interface. Both helices have been shown to be foldons which initiate the folding process of cytochrome c [8]. Other parts of the structure are primarily composed by coil regions, fixate a heme ligand, and show low SR. (**b**) Per residue SR as line chart. The standard deviation is given for each point. Residues without contacts exhibit a relevance of 0 Å.

The SR score computed by StructureDistiller of many residues of cytochrome c is neutral or even negative. Especially coil regions feature contacts which tend to decrease reconstruction fidelity. Remarkable are the high SR scores of HIS-26 and GLY-45 as well as their direct contact for which the score amounts to 0.172 Å (making it the fifth most relevant contact). No SR is reported for PRO-44 as it does not participate in any contacts according to the employed contact definition, though both groups are positioned in a way which would allow them to form a hydrogen bond. In literature [52], the contact between HIS-26 and PRO-44 is reported as crucial for the correct conformation of cytochrome c. Disruptions will result in a loss of structure [52], though the relevance of PRO-44 may also be attributed to the backbone rigidity introduced by the proline residue. The detection of relevant contacts and positions is fuzzy [54], but the high scoring contact between HIS-26 and GLY-45 implies the importance of a contact between both Ω-loops for successful protein folding as well as structure reconstruction. Between GLY-29 and MET-80 the most relevant contact (with the highest SR) is located, it increases reconstruction fidelity by 0.563 Å on average. This contact occurs between two unordered coil regions as well and implying that some structural information on the correct arrangement of these unordered protein parts is crucial for a successful reconstruction. Mutations to HIS-33 have been demonstrated to show no effect [52] which is also captured by slightly negative SR score of −0.015 Å. Both N- and C-terminal helix contain residues with high relevance, especially in regions where both helices interact. The importance of these helix contacts has been shown previously [55]. The role of both helices as foldons [8] points to a high intrinsic stability.

The SR score successfully spots contacts and residues crucial for structure integrity as shown in experiments [52, 8, 55]. The previously described contact between HIS-26 and PRO-44 [52] is absent as the result of a too strict contact definition, yet the necessity of structural information in this region is captured nevertheless.

## Discussion

Contact maps are one of the most prominent tools in today’s structural bioinformatics [29, 19], though mere knowledge of residue contacts can neither describe all events of the protein folding process [56] nor is it the optimal basis of structure prediction techniques [38]. Our study demonstrates that native contacts in a protein structure are not of equal importance for the reconstruction of the tertiary structure from this reduced representation. A more fine-grained interpretation of contact maps can be achieved by the consideration of our proposed Structural Relevance scores. Contacts of high Structural Relevance tend to be unique contacts for which no redundant backup exists as it is the case for the contact between two Ω-loops in cytochrome c [52]. The importance of this contact for the structural integrity also implies that high Structural Relevance scores may capture crucial positions for structure stability.

Coevolution or supervised machine learning techniques are the basis for the prediction of contact maps [49, 29, 21]. Conventionally, contact predictors are designed and trained on collections of all native contacts in a dataset. Subsequently, the most reliable contacts are selected from all predictions; the size of this subset depends on sequence length [35]. This study shows that these subsets drastically change in meaningfulness as indicated by reconstruction fidelity. An implication is that it is not the optimal strategy to consider a random subset of contacts; reconstruction fidelity and information per contact will be increased when the contacts with the highest Structural Relevance scores are considered. Especially, coevolution techniques struggle with transitive predictions: if there is a signal between residue A and B as well as B and C, then likely for the pair A and C a signal is reported too [18, 49]. Machine learning also comes at a price: an increase of true positive predictions involves more false positives. Furthermore, focusing on the most relevant contacts may make it easier to spot the driving forces of protein folding. Contact maps and reconstruction algorithms can compensate some false positive predictions [23], but errors in prediction methods tend to be correlated and hinder reconstruction more than random noise [36]. Regardless of the particular approach, it may be beneficial when predictors do not try to cover all native contacts but focus only on the most relevant ones. This would decrease the number of predicted contacts but may increase the reliability of their prediction by avoiding both false positive predictions and emphasizing contacts which promise to improve reconstruction fidelity the most while ignoring those which contribute only marginally.

Residues in a protein are covalently bound and constraints on a residue will also affect neighboring residues. Thus, residue-specific information as complex as the Structural Relevance score should not be considered the absolute truth [54]. One of the most delicate aspects when handling contact maps is the used contact definition [24, 38]. Particularly, the distance-based contact definition employed in this study does not imply chemically relevant contacts between atoms (such as hydrogen bonds or hydrophobic interactions). The chosen cutoff is rather strict and will ignore some meaningful contacts; a relaxation of this cutoff will encompass more contacts but also increases computation time. In some cases such as the assembly of helices [57], information of the complex hydrogen bond patterns has to be considered, which may benefit from a more fine-grained contact definition. Another refinement of the proposed protocol is the explicit consideration of residues which are not in contact. This information has been demonstrated to increase reconstruction fidelity [38]. Also, it is natural that the employed reconstruction pipeline [22] as well as scoring scheme [58] have an effect on the computed scores and may introduce some form of bias. The TM-score may be more suited to score reconstructs because it is independent of protein length and can more intuitively state whether the correct protein fold was reconstructed [59]. TM-score values are provided as output, but presented results use the RMSD value because of its widespread use and comparable results in relation to previous studies [40, 41, 24]. Furthermore, the decision to use 30% of all native contacts to compute the Structural Relevance score may be not generally applicable. The StructureDistiller algorithm may be improved by determining for each protein structure individually where the sweet spot lies between meaningful reconstructs and maximized sensitivity.

In summary, the StructureDistiller algorithm is presented as an approach to assess the structural relevance of individual contacts and residues. This constitutes a novel contribution of the toolkit available for the interpretation of contact maps and protein structures in general, while making the connection of contact maps and tertiary structure more concrete. Results of the StructureDistiller algorithm may enhance contact prediction techniques [18, 19, 20, 21], contact map evaluation [35, 37], reconstruction algorithms [22], quality assessment programs for the yielded models [60], statistical potentials, and may even give particularized insights into the protein folding problem itself [41].

On a general level, the dataset of Early Folding and Highly Stable Residues [10] provides valuable information to converge on the protein folding problem [9, 3]. The Start2Fold dataset [10] enables the direct connection of protein folding and structure prediction which is furnished by contact map representations. It is implied that Early Folding Residues may initiate protein folding and determine the order in which local structures are assembled [2, 12] but they are of average relevance in terms of the Structural Relevance score. Highly Stable Residues may not fold early but constitute regions of a protein which prevent spontaneous unfolding. Interestingly, regions of Highly Stable Residues are of high relevance for the formation and stabilization of the correct protein fold. David Baker [61] showed that short-range contacts lead to fast folding whereas a high ratio of long-range contacts leads to a slow down. Early Folding Residues initiate the folding process by establishing contacts to neighbors at sequence level [2, 44]. Furthermore, hydrophobic interactions, contacts of ordered secondary structure elements, as well as long-range contacts promote structural integrity. In a previous study [5], we showed that Early Folding Residues are embedded in ordered secondary structures and are embedded in a network of hydrophobic interactions. This may imply that Early Folding Residues may initiate the formation of local structures which can then assemble to actually stabilize the global structure of a protein by Highly Stable Residues.

Maybe the protein folding problem is not solvable without understanding how protein structures can be predicted reliably. Indeed, both problems are often described to be two sides of the same coin [16] and structure prediction did provide new insights into the folding process before [42, 43]. Additional tools are needed to make the connection of protein sequence and structure more tangible and StructureDistiller provides just that. The algorithm allows for a novel fine-grained interpretation of contact maps and may improve their interpretability. Applications of the proposed algorithm are not limited the Start2Fold database [10], it can be used for the analysis of arbitrary protein structures, e.g. to assess structural effects of mutations at certain residue positions. Following this new paradigm, the interface between protein folding and structure prediction [16] can be explored in more detail.

## Methods

### Datasets used for evaluation

The Start2Fold database [10] provides results of pulse labeling hydrogen-deuterium exchange experiments. For the 30 proteins of the dataset (see Supplementary Figure 1 and supplementary material of [3] for a detailed definition), 5,173 contacts of 2,529 residues were evaluated. Positions without native contacts were ignored. The Start2Fold database was chosen because it provides a standardized annotation of EFR which initiate the folding process [3, 44, 4] and HSR which exhibit significant resilience to unfolding events [10]. This dataset encompasses all major CATH and SCOP classes. Thus, the SR score was assessed using a dataset of proteins for which the folding characteristics are fairly well understood. The size of proteins in the dataset varies from 56–164, which emphasizes relatively small proteins. The covered fold classes are diverse, but present proteins tend to be single domain proteins with fast folding kinetics [44]. Entries without EFR annotation were ignored, even when information on HSR was present. Residues were considered buried when their relative accessible surface area was below 0.16 [62]. Evolutionary couplings were computed by the EVfold web server [18, 49]. BioJava [63, 64] implementations of the algorithm of Shrake and Rupley [65] and DSSP [66] were used for accessible surface area and secondary structure element computation respectively.

### Annotation of residue contacts

A pair of residues was defined to be in contact when the distance between their *C*_*α*_ atoms was less than 8 Å. Contacts maps were created based on this contact definition while ignoring contacts between residues less than six positions apart at sequence level. The remaining tertiary contacts were considered short (sequence separation of 6–11), medium (12–23), or long (>23) [39]. Non-covalent interactions (i.e. hydrogen bonds and hydrophobic interactions) were annotated by PLIP [46].

### Structure reconstruction and performance scoring

Contact maps (or subsets thereof) were reconstructed as all-atom models by CONFOLD [22]. Secondary structure information was annotated by DSSP [66] and provided as input of the reconstruction routine. By default, CONFOLD creates a set of reconstructs and selects the five top-scoring ones as output. The selected reconstructs and the native structure were then superimposed and their dissimilarity was measured by the RMSD. TMalign [58] was used for alignment.

### The StructureDistiller algorithm

The StructureDistiller algorithm (Figure 7) evaluates the structural relevance of individual contacts in the context of a set of other contacts. By selecting 30% of the native contacts of a map, baseline reconstructs can be created which resemble the protein fold and are highly sensitive to the toggling (removal or addition) of an individual contact. The performance of the baseline reconstructs can be quantified by a structural alignment to the native structure. Analogously, the performance can be measured for the toggle reconstructs, which represent the change of one particular contact. By comparing the performance of a toggle reconstruct with its corresponding baseline reconstruct, the SR score of all contacts is quantified.

**Figure 7:**
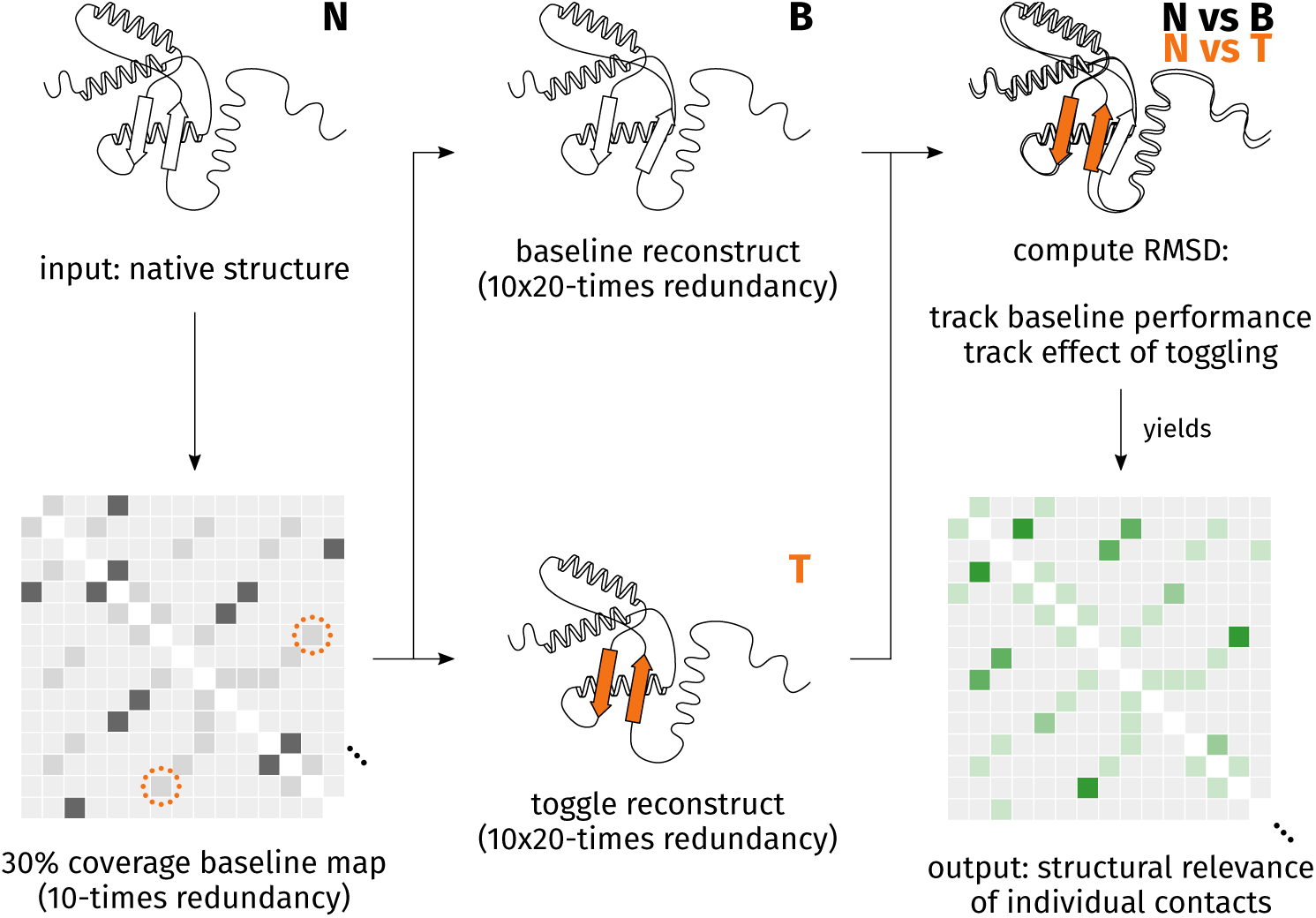
Depiction of the StructureDistiller algorithm. In order to compute the SR score of individual contacts, the effect of their consideration on the reconstruction performance (ΔRMSD) is measured. This allows a novel, more fine-grained interpretation of contact maps. By using 30% of all contacts present in the native structure (N), baseline contact maps are created which provide maximum sensitivity to the removal or addition of a single contact. Baseline reconstructs (B) provide the context to assess the role of individual contacts. Within each baseline contact maps, all contacts of the native contact map are toggled: contacts already present are removed and those absent are added (depicted by orange circle of dots). Reconstructs are created based on these toggle contact maps (T). By superimposing reconstruct and native structure, the SR score of all contacts can be quantified as relative change in RMSD. The idea is that some contacts may provide information which is crucial for reconstruction fidelity, e.g. on the correct arrangement of secondary structure elements (depicted by orange fill).

The StructureDistiller algorithm is represented in Algorithm 1. A protein structure *S*_native_ in legacy PDB format is the input. Structure files should encompass single domains of a single chain. The corresponding contact map *C*_native_ is created. *C*_native_ constitutes the set of all contacts which will be evaluated.

Fractions equal to 30% of *C*_native_ are then randomly selected. The SR score of a contact depends on all other contacts used for a reconstruction. No effect can be expected when a contact is considered which contributes no additional, but only redundant information [41]. The creation of random subsets of *C*_native_ is performed with a redundancy *r* of 10. The resulting subset of contacts *C*_baseline,*i*_ is used to create the baseline reconstructs *S*_baseline,*i*_. The average RMSD_baseline,*i*_ of each created subset *C*_baseline,*i*_ is tracked with respect to *S*_native_. These subsets are highly sensitive to the removal and addition of a single contact and the basis for all further computations.

All contacts of *C*_native_ are now evaluated regarding their SR by pairing each contact to each baseline subset of contacts *C*_baseline,*i*_. For each pair, it is determined whether the current contact *c* is element of *C*_baseline,*i*_. If so, *c* is removed from *C*_baseline,*i*_, else *c* is added to the corresponding subset. The change in reconstruction performance can be quantified by this toggling of a contact: the modified subset *C*_toggle,*i*_ is again used for a reconstruction and RMSD_toggle,*i*_ is used to describe its quality. The average improvement of the reconstruction with knowledge of the contact *c* is tracked by ΔRMSD_*c*_. RMSD_baseline,*i*_ − RMSD_toggle,*i*_ is evaluated when *c* was removed from the subset, the expression is flipped when *c* was added. The SR of individual residues is the average of all ΔRMSD_*c*_ of contacts this residue participates in. Positive SR scores represent contacts which increase reconstruction fidelity while negative scores occur for contacts hindering reconstruction.

The runtime of StructureDistiller scales with the number of contacts in the initially created map *C*_native_. The individual reconstruction tasks are distributed among worker threads which allows for efficient parallelization. Using a conventional workstation, computation on proteins with up to 200 residues requires one day on average.

### Definition of reconstruction strategies

Various subset selection strategies were used to assess the relevance of contacts in a contact map. In all cases, a number equal to 30% of the contact count in the native map was used. For the creation of the random bin, 30% of all native contacts were chosen randomly. The most relevant selection constitutes the 30% of all contacts sorted for highest SR, least relevant resembles 30% of all contacts with the lowest scores. All percentage numbers are relative to the number of contacts in the native structure. All operations on all definitions are performed with ten-fold redundancy. Contact distances were assessed: all short (sequence separation of 6–11) and long (>23) contacts [39] were assessed. The same was done for hydrogen bonds and hydrophobic interactions. Because the number of contacts of a particular distance or type may be smaller than 30%, a dedicated bin (e.g. non-short) was created to match in size.

#### Algorithm 1 StructureDistiller Pseudocode

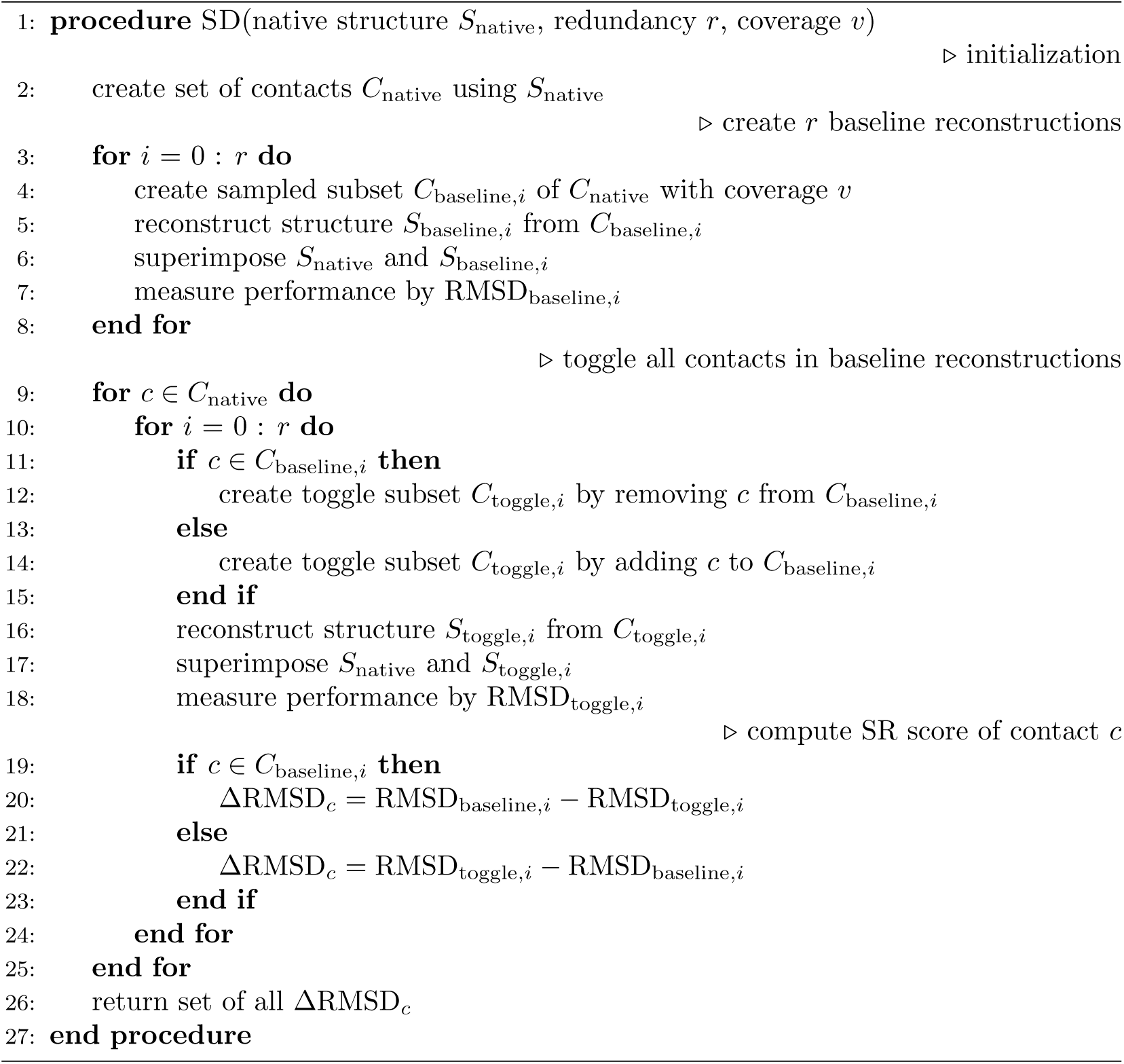

### Introduction of false positive contacts

False positive contacts are contacts not present in the contact map of the native protein structure. Contact maps were created by the best and random strategy and in 1% bins up to 10% false positive contacts were introduced, replacing the initially selected native contacts. Analogous to the employed contact definition, false positive contacts were required to exhibit a sequence separation greater than five.

### Statistical analysis

Residues without any contacts (i.e. where no SR score can be computed) were ignored from statistical analysis. Notched box plots were used for visualization. The notch corresponds to the 95% confidence interval around the median. When the notches of two distributions do not overlap, they can be assumed to be different. Significance was explicitly tested by a two-tailed Mann-Whitney U test. *p*-values <0.05 were considered significant.

## Data availability

A reference implementation of the StructureDistiller algorithm is available in the module structural-information at https://github.com/JonStargaryen/jstructure. A compiled version is deposited at https://doi.org/10.5281/zenodo.1405369. All evaluated data is included in the manuscript and its supplementary information.

## Acknowledgements

The authors thank Jose Duarte, Christoph Leberecht, Sarah Krautwurst, and Florian Kaiser for scientific discussions and/or proofreading of the manuscript. Support for S.B. within the RCSB PDB comes from the National Science Foundation, the National Institutes of Health, and the Department of Energy (NSF DBI-1338415; Principal Investigator: Stephen K. Burley).

## Author contribution

S.B. conceived the study, implemented the algorithm, and analyzed the data. M.S. and D.L. supervised the research. All authors revised the manuscript.

## Competing interests

None declared.

