## Supplemental Material for "StructureDistiller: Structural relevance scoring increases resilience of contact maps to false positive predictions"

**Supplementary Figure 1**

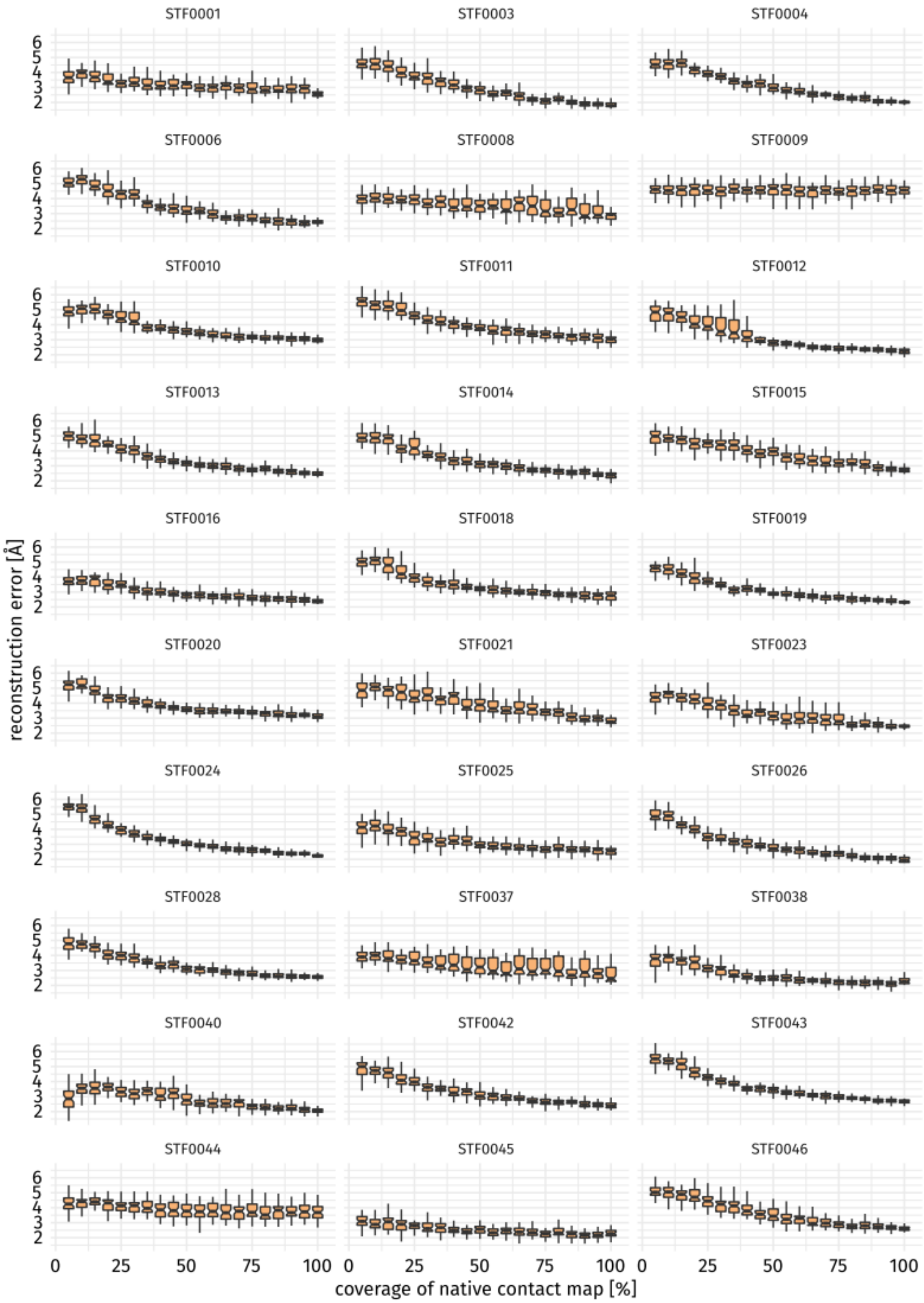


**Reconstruction error by percentage of contacts split by protein.**
For some proteins the reconstruction performance does not increase when more or even all native contacts are considered. In other cases, reconstructions with a large number of constraints exhibit high variance regarding their reconstruction performance.

**Supplementary Figure 2**

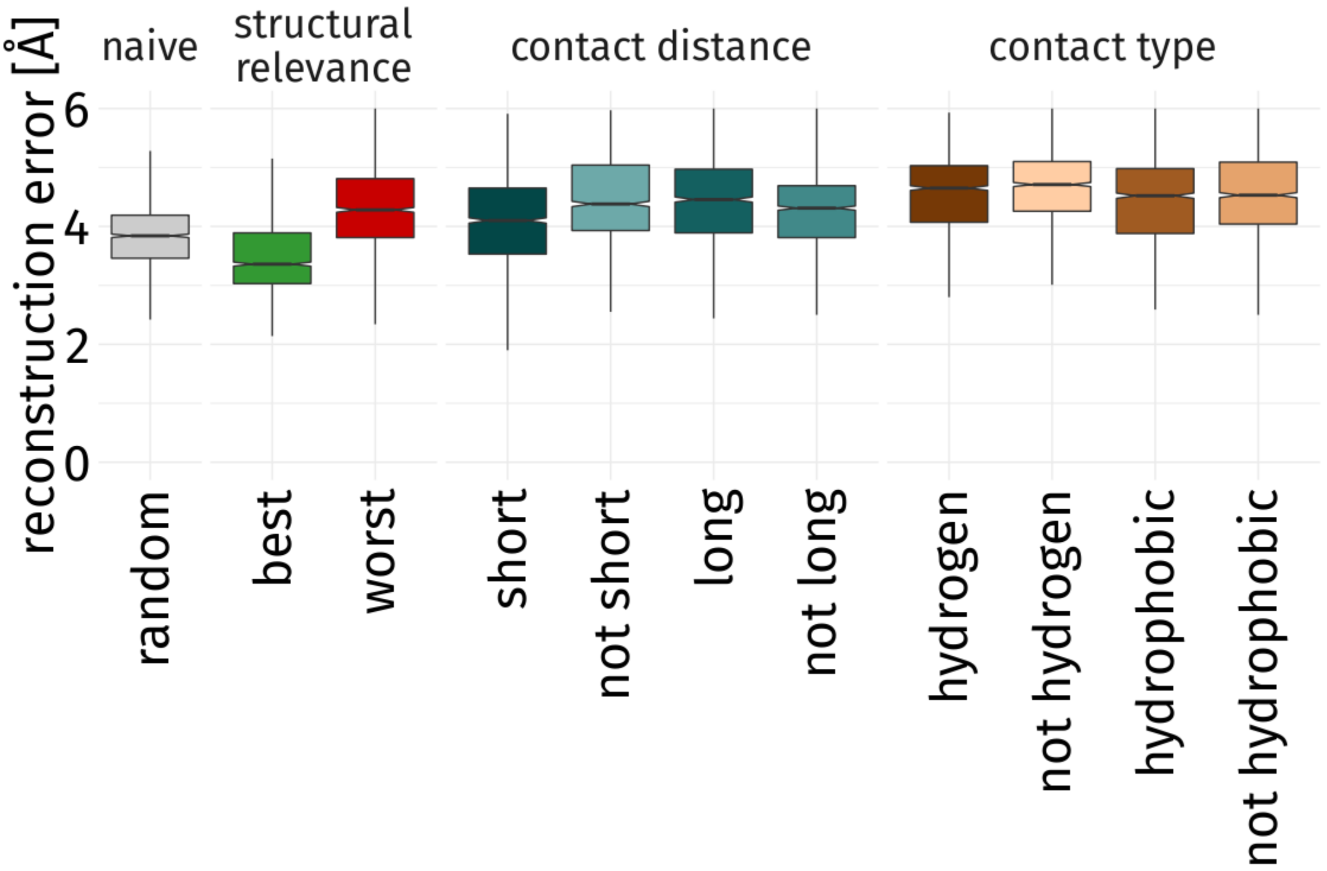


**Detailed impact on reconstruction performance by strategy.**
Various strategies were used to reconstruct structures of the dataset using a number of constraints equal to 30% of contacts in the native map. Contact distance and type bins are only comparable to the explicitly negated bins because the available number of contacts differs (i.e. there may not be enough hydrogen bonds to match the number of contacts in the random bin). Not selections based on contact distance or type performe worse or on par than their counterpart which implies the necessity to consider a complex collection of contacts for a successful reconstruction (see Chen et al., 2007). When combined short contacts yield relatively good reconstructs even though they structural relevance scores are low.
